# Growing Macrophages Regulate High Rates of Solute Flux by Pinocytosis

**DOI:** 10.1101/2024.10.22.619691

**Authors:** Biniam M. Tebeje, Natalie W. Thiex, Joel A. Swanson

**Affiliations:** Department of Microbiology and Immunology, University of Michigan Medical School, Ann Arbor, MI 48109-5620; Department of Biology and Microbiology, South Dakota State University, Brookings, SD 57007

## Abstract

In metazoan cells, growth factors stimulate solute ingestion by pinocytosis. To examine the role of pinocytosis in cell growth, this study measured cell proliferation and the attendant rates of solute flux by pinocytosis in murine macrophages in response to the growth factor colony-stimulating factor-1 (CSF1). During CSF1-dependent growth in rich medium, macrophages internalized 72 percent of their cell volume in extracellular fluid every hour. Removal of the essential amino acid leucine from growth medium limited rates of protein synthesis and growth, but increased rates of solute accumulation by macropinocytosis. The amount of protein synthesized during leucine-dependent growth exceeded the capacity of pinocytosis to internalize enough soluble leucine to support growth and proliferation. Fluid-phase solute recycling from lysosomes secreted small molecules from the cells at high rates. Inhibitors of pinocytosis and the mechanistic target-of-rapamycin (mTOR) reduced cell growth and solute recycling, indicating roles for pinocytosis in growth and for nutrient sensing in the regulation of solute flux by pinocytosis.

**Summary:** Murine macrophages growing in response to colony-stimulating factor-1 (CSF1) require pinocytosis. High rates of solute influx and accumulation by pinocytosis are regulated by CSF1 and leucine. Low molecular weight products of protein hydrolysis in lysosomes recycle efficiently from the cells.

## Introduction

In pinocytosis, cells internalize extracellular fluid and solutes by invagination of plasma membrane into intracellular vesicles called pinosomes. Small pinosomes form by various kinds of coated vesicles, such as clathrin-coated vesicles. Larger pinosomes, called macropinosomes, form by actin-mediated motile activities (Mylvaganam et al., 2021; Swanson, 2008). Newly formed pinosomes recycle membrane and solutes to the cell surface and the extracellular space, fuse with other pinosomes or endosomes, and fuse with lysosomes, where internalized solutes accumulate or are degraded by acid hydrolases (Kay, 2021). Water, ions and small molecules such as amino acids cross membranes of pinosomes or lysosomes into cytoplasm (Chadwick et al., 2021; Hu et al., 2022).

The growth of metazoan cells is regulated by growth factors, which are secreted proteins that promote cell proliferation by binding to receptors on cell membranes and activating biochemical reactions that support anabolic metabolism, including protein synthesis (Thompson and Bielska, 2019). An association between growth factor signaling and increased pinocytosis has been known for decades (Brunk et al., 1976; Davies and Ross, 1978; Haigler et al., 1979; Racoosin and Swanson, 1989). Yet to the best of our knowledge no studies have analyzed the role of growth factor-stimulated pinocytosis in cell growth.

In macrophages, growth and differentiation are regulated by the growth factor colony-stimulating factor-1 (CSF1, or macrophage colony-stimulating factor, M-CSF) (Stanley and Chitu, 2014). CSF1 stimulates macrophage growth and macropinocytosis in a dose-dependent manner (Lou et al., 2014; Racoosin and Swanson, 1989; Tushinski et al., 1982). CSF1 binding to its cognate CSF1 receptor in plasma membranes initiates macropinocytosis as well as many intracellular signals necessary for growth, which include activation of class 1 phosphatidyl-inositol 3-kinase (PI3K-1), Akt, and the mechanistic Target-of-Rapamycin Complex-1 (mTORC1) (Stanley and Chitu, 2014; Yoshida et al., 2018).

In addition to its association with growth factor signaling, macropinocytosis is associated with growth itself. Increased macropinocytosis of extracellular proteins by Ras-transformed cancer cells permits their growth in otherwise nutrient-poor tumor environments (Commisso et al., 2013; Kamphorst et al., 2015). Furthermore, macropinocytosis in the amoeba *Dictyostelium discoideum* allows it to grow in axenic media (Kay et al., 2019).

Macropinosome formation in response to CSF1 requires PI3K-1 (Araki et al., 1996), and growth factor-dependent activation of mTORC1 in macrophages and fibroblasts requires internalization of the essential amino acid (EAA) leucine into macropinosomes (Yoshida et al., 2015). Activation of mTORC1 in T cells requires ingestion of EAA into macropinosomes (Charpentier et al., 2020). These studies suggest that growth factor signaling for cell growth requires ingestion of amino acids by macropinocytosis. However, many cells acquire EAA through transporters in plasma membrane (Gauthier-Coles et al., 2021; Nicklin et al., 2009), potentially obviating any need for pinocytosis in amino acid entry. Thus, although activation of mTORC1 for increased anabolic metabolism requires macropinocytosis of leucine, it is not known whether growth factor-stimulated macropinocytosis can ingest enough amino acids to support the protein synthesis needed for growth and proliferation.

Another indication that pinocytosis may be essential for cell growth is that it is regulated by the availability of amino acids, particularly EAA (Besterman et al., 1983). Cells grown in protein-rich media deficient in EAA can upregulate macropinocytosis, which supports cell growth by scavenging of extracellular proteins, mostly albumin (King et al., 2020; Nofal et al., 2022).

Conversely, incubation of macrophages in leucine and other EAAs inhibits macropinocytosis in response to acute stimulation by CSF1, by initiating the loss of CSF1R from cell surfaces (Mendel et al., 2022). Thus, rates of solute accumulation by macropinocytosis can be regulated by nutrient availability.

These many correlations prompt questions about the role of growth factor-stimulated pinocytosis in cell growth. What is the magnitude of solute flux through growing cells and is that flux sufficient to support growth by ingestion of nutrients? We hypothesize that CSF1-stimulated pinocytosis is necessary for macrophage proliferation.

Pinocytosis in macrophages can be reliably measured using Lucifer yellow (LY), a membrane-impermeant, non-degradable, small fluorescent dye (Stewart, 1978). Accumulation of LY by macrophages is proportional to its concentration in medium and is inhibited by metabolic inhibitors and by incubation at 4 C (Swanson et al., 1985). LY accumulation by macrophages is the net result of high rates of influx and efflux by vesicular traffic to and from lysosomes. Influx can be measured in the earliest rates of LY accumulation (1-5 minutes), before significant recycling occurs. Efflux can be measured as the loss of LY from cells which have been preloaded by pinocytosis, washed, and chased in unlabeled medium. The kinetics of LY efflux indicate two compartments that recycle solutes exponentially: a rapidly emptying compartment, likely pinosomes or early endosomes, and a slowly emptying compartment, likely lysosomes (Swanson et al., 1985).

To analyze solute flux as a component activity of cell growth, this study quantified pinocytosis in growing primary murine macrophages and measured the effects of pinocytosis inhibitors on CSF1-dependent growth and proliferation. Using LY to measure pinocytosis, we found that growing macrophages maintained high rates of solute influx and efflux. Using leucine-limited growth media, we report that the volume of medium ingested by pinocytosis is not sufficient to support leucine-dependent growth. The high rates of solute efflux indicated that small molecules generated by hydrolysis in lysosomes are excreted from growing macrophages. Moreover, inhibitors of pinocytosis limited cell growth, and the mTORC1 inhibitors rapamycin and Torin-1 increased net solute accumulation, indicating a role for mTORC1 in the regulation of pinocytosis in macrophages. These studies indicate high rates of water and solute flux into and through growing cells and establish pinocytosis as essential to growth factor-dependent proliferation of macrophages.

## Results

### Growing macrophages are actively macropinocytic

Growth, proliferation and pinocytosis by cultured primary bone marrow-derived macrophages were measured in a rich growth medium (bone marrow medium, BMM). Growth and proliferation of macrophages were quantified using an Incucyte SX1 system, which regularly collected multiple phase contrast images of cells from individual wells of 24-well tissue culture plates growing in a 37 C, CO_2_ incubator. Images of macrophages in BMM, with and without 50 ng/ml CSF1, were recorded every hour for 1-48 hrs (Fig. 1A). Image analysis software that measured cell numbers per well as a function of time showed CSF1-dependent proliferation (Fig. 1B), which was confirmed by measuring total protein content of the wells in parallel growth studies (Fig. 1C). Protein assays of lysates from cells plated at different densities and counted in the Incucyte determined that protein content of macrophages growing in BMM was 93 pg protein/cell (Fig. 1D). Growth-associated protein synthesis was then quantified at each time point as cell number times the average protein content per cell. In BMM with CSF1, macrophages grew at a rate of 31,800 cells_*_10^6^ cells^-1^ hr^-1^, or 2.96 mg protein_*_10^6^ cells^-1^ hr^-1^.

**Figure 1.**
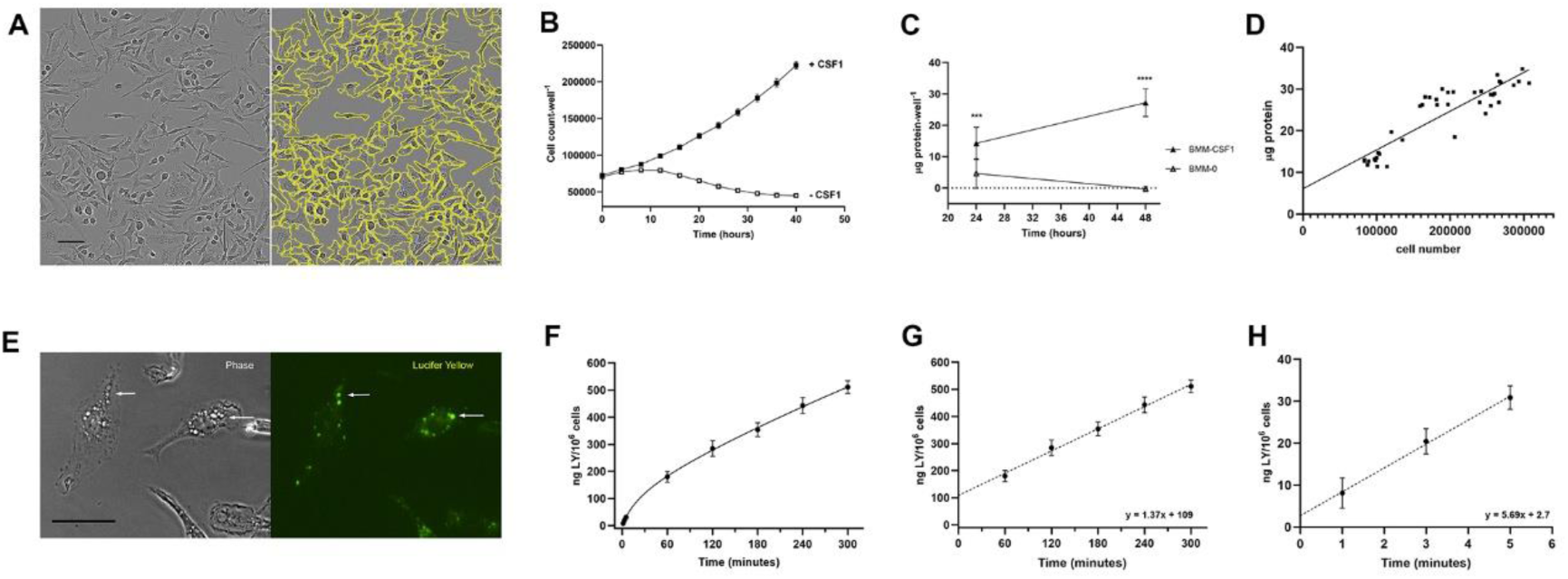
Macrophages growing in rich medium are highly pinocytic. (A) Images of macrophages growing in BMM. Left image shows phase contrast image of macrophages in a 24-well tissue culture plate in the Incucyte SX1. Right image shows calculated cell outlines (yellow) used to determine cell numbers per well. Scale bar: 50 mm. (B) Cell numbers per well of macrophages growing in BMM, +/- CSF1, as measured hourly. Measurements from every 4 hrs are shown. Data show mean and SEM obtained from 9 wells per time point (3 biological replicates). (C) Protein accumulation by cells growing in BMM, +/- CSF1, measured from cell lysates collected at 24 and 48 hrs. Differences measured at each time point with and without CSF1 were significant (***, p<0.001; ****, p<0.0001). Data show mean and SEM of protein_*_well^-1^, obtained from 9 wells per condition (3 biological replicates). (D) Measurement of protein content per cell in BMM. The plot shows the relationship between cell number per well and mg protein per well (3 biological replicates). The mean protein content was 93 pg*cell^-1^ (linear regression: mg protein/well = (.000093 mg protein/cell x cell number/well) + 6.1 mg protein/well, R^2^ = 0.79). (E) Macropinosomes in macrophages in CSF1. Macrophages on coverslips (MatTek dishes) were incubated in 500 mg_*_ml^-1^ LY in BMM/CSF1 for 5 min, then were washed free of LY and fixed for microscopic observation of phase contrast (left) and LY fluorescence (right). Scale bar: 20 mm. (F) Time course of LY accumulation by cells growing in BMM. Cells were lysed after 1, 3, 5, 60, 120, 180, 240 and 300 min in BMM/CSF1 with 500 mg_*_ml^-1^ LY before washing and lysis. Fluorescence and protein content of each well were measured. Protein was converted to cell number to obtain ng LY per 10^6^ cells. Data show mean and SEM from 9 wells per time point (3 biological replicates). (G) Rate of LY accumulation. Linear regression of data in panel F, 60-300 minutes. Slope = 1.37 ng LY_*_10^6^ cells^-1^ min^-1^, b=109, R^2^=0.95 (= 82 ng LY_*_10^6^ cells ^-1^ hr^-1^, = 164 fl fluid_*_cell^-1^ hr^-1^). (H) Rate of influx. Linear regression of data in panel F, 1-5 minutes (influx). 5.7 ng LY_*_10^6^ cells^-1^ min^-1^, b=2.7, R^2^=0.90 (= 342 ng LY 10^6^ cells ^-1^ hr^-1^, = 683 fl fluid cell^-1^ hr^-1^).

To ask whether growing macrophages ingest enough nutrients by pinocytosis to support these growth rates, pinocytosis was measured using LY. Incubating cells for 5 minutes in medium containing LY, followed by fixation and microscopic observation, showed that growing macrophages contained many macropinosomes (Fig. 1E). To measure pinocytosis of LY in populations of cells, macrophages in 24-well dishes were incubated for various times in BMM containing 500 mg_*_ml^-1^ LY, then dishes were emptied and washed by immersion into cold phosphate-buffered saline (PBS). Cells were lysed in 0.1 % Triton X-100 and the fluorescence and protein content of each well was measured. Protein measurements were converted to cell numbers per well to obtain LY accumulation per 10^6^ cells. As in previous studies (Berthiaume et al., 1995; Swanson et al., 1985), accumulation was curvilinear initially, then linear after 30-60 minutes (Fig. 1F, Suppl. Fig. 1A). Linear regression analysis of LY accumulation from 60-300 min indicated that macrophages accumulated 1.37 ng LY_*_10^6^ cells^-1^ min^-1^ or 82 ng LY_*_10^6^ cells^-1^ hr^-1^ (Fig. 1G). Correcting for the concentration of LY in medium determined that growing macrophages accumulated 164 femtoliters (fl) of fluid per cell per hour.

Previous studies of pinocytosis showed that solute accumulation is the net result of solute influx and efflux (Besterman et al., 1981; Swanson et al., 1985). Influx reflects the rate of pinosome formation, which can be measured in the early rates of LY accumulation (1-5 minutes). In macrophages growing in BMM, the LY influx rate was 5.7 ng LY_*_10^6^ cells^-1^ min^-1^ or 342 ng LY_*_10^6^ cells^-1^ hr^-1^, 4.2 x greater than the rate of net accumulation (Fig. 1H). This rate indicates that a macrophage growing in 50 ng_*_ml^-1^ CSF1 ingests 684 fl of extracellular fluid per hour and recycles a large portion of the ingested solutes. Using earlier size measurements of CSF1-cultured murine macrophages (12.2 mm diameter) (Falk and Vogel, 1988), and calculating their volumes as spheres of that diameter, we estimate their volume at 951 mm^3^ (951 fl). Therefore, growing macrophages internalize the equivalent of 72% of their cell volume in extracellular fluid every hour. Given this high rate of influx and the concentrations of amino acids and proteins in BMM, this suggests that if all amino acids internalized into pinosomes could be imported across macropinosome or lysosome membranes into cytoplasm, then macrophages could potentially synthesize the protein required for cell growth using the nutrients taken in by pinocytosis alone.

### Leucine-dependent growth occurs at low concentrations of leucine

To examine whether pinocytosis can supply macrophages with enough free amino acids for growth, we measured CSF1-dependent growth and pinocytosis in leucine-limited conditions. As an essential amino acid, leucine in macrophages must be obtained from exogenous sources. We prepared RPMI medium containing dialyzed fetal bovine serum (dFBS) but lacking added leucine, which we refer as R(0). Macrophages cultured in R(0) would have access to some leucine derived from lysosomal degradation of proteins in dFBS, so we expected some cell growth in R(0) with CSF1. But compared to non-dialyzed FBS, dFBS lacks amino acids, peptides and metabolites smaller than 10,000 Da. R(0) was supplemented with 800 mM leucine (R(Leu), the concentration of leucine in RPMI), 800 mM isoleucine (R(Iso), still lacking leucine), or 10 mg_*_ml^-1^ bovine serum albumin (BSA; R(BSA)) and with 50 ng_*_ml^-1^ CSF1 (R(0)/CSF1, R(Leu)/CSF1, R(Iso)/CSF1, R(BSA)/CSF1). Macrophages died without CSF1 (Fig. 2A). In CSF1-containing media, growth was initially unaffected by the presence or absence of leucine, but leucine-dependent proliferation appeared after 15 hrs. Proliferation was measurable in all the media but was greater in R(Leu)/CSF1 and R(BSA)/CSF1 than in R(0)/CSF1 (Fig. 2A). Growth was measured as the rate of change in cell number from 30-38 hr of culture in the various media, normalized to the cell number at 30 hr (Fig. 2B). Subtracting the rate of growth in R(0)/CSF1 (5800 cells_*_10^6^ cells^-1^ hr^-1^) from the rate of growth in R(Leu)/CSF1 (26,100 cells_*_10^6^ cells^-1^ hr^-1^) or in R(BSA) (19,000 cells 10^6^ cells^-1^ hr^-1^) yielded leucine-specific growth rates of 20,300 cells_*_10^6^ cells^-1^ hr^-1^ in R(Leu) /CSF1 and 13,200 cells_*_10^6^ cells^-1^ hr^-1^ from the leucine potentially available from lysosomal degradation of the added protein in R(BSA) (Fig. 2C). Growth rates in R(Iso) were significantly less than those in R(Leu)/CSF1, indicating the specificity of growth for leucine (Suppl. Fig. 2A).

**Figure 2.**
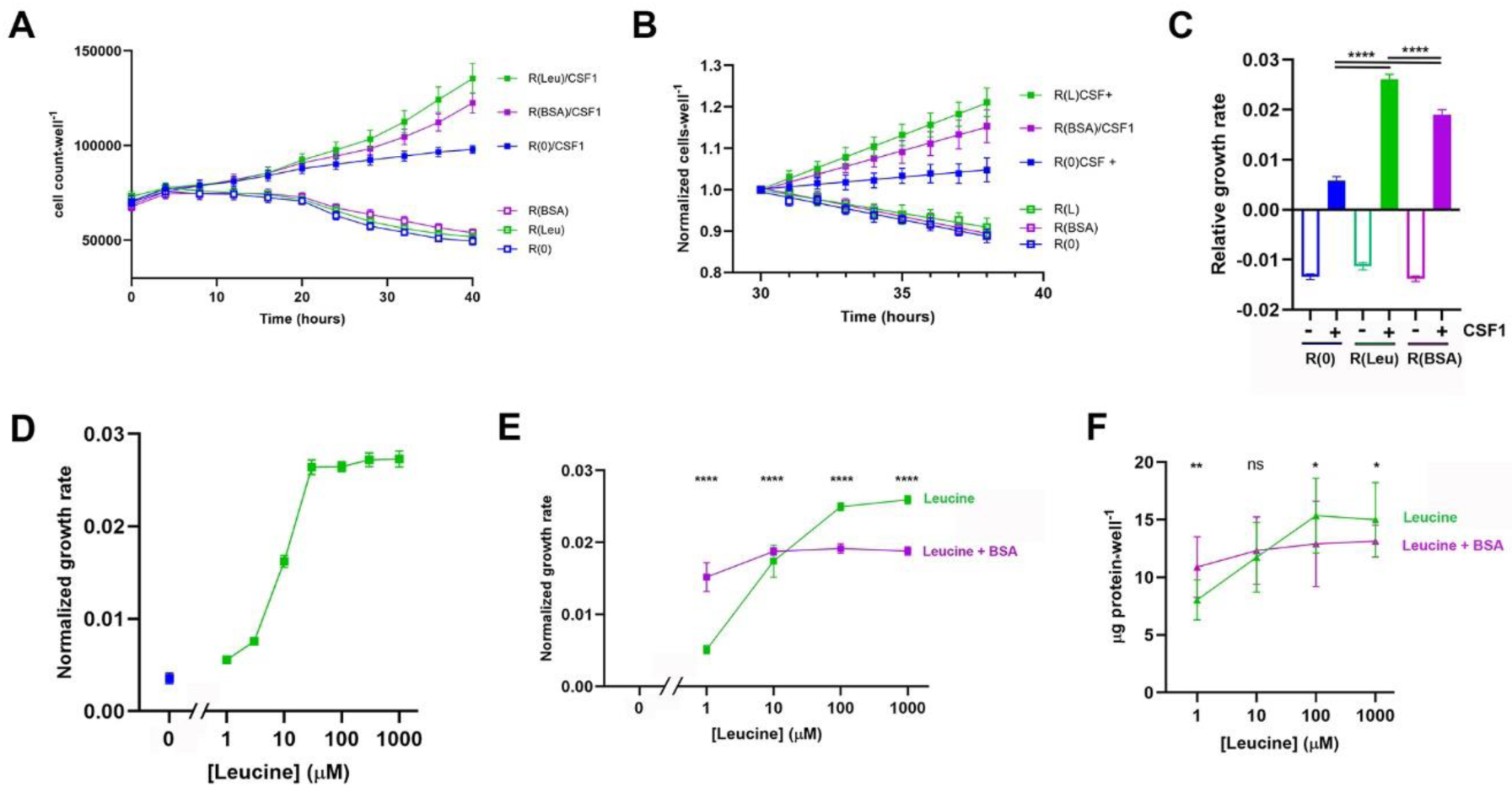
Leucine-dependent growth occurs at low leucine concentrations and is modulated by BSA. (A) Time course of cell growth in R(0), R(Leu), and R(BSA), +/- CSF1, as measured hourly. Measurements from every 4 hrs are shown. Data show mean and SEM obtained from 9 wells per time point (3 biological replicates, 3 wells per condition). (B) Rates of cell growth in R(0), R(Leu), R(BSA), +/- CSF1 from the time courses in panel A, showing hourly measurements normalized to the cell counts for each condition at 30 hrs. Lines show linear regression analysis (n = 81 per condition). (C) Slopes of the lines shown in panel B. The relative rates of cells in media with CSF1 were highly significant (One-way ANOVA, ****, p<0.0001). (D) Rates of cell proliferation in R(0) (blue) and in R(0) supplemented with the indicated concentrations of leucine (green). Cell numbers per well were measured every hour and normalized to the cell numbers per well at 30 hrs (as in panel 1B). Each point indicates the mean and SEM of a slope determined from 81 measurements (9 time points from 30-38 hrs, 3 wells per time point, 3 biological replicates). (E) Effects of BSA (10 mg_*_ml^-1^) on rates of cell proliferation in different concentrations of leucine. Each point indicates the mean and SEM of a slope determined from 81 measurements (9 time points from 30-38 hrs, 3 wells per time point, 3 biological replicates). BSA stimulated growth in low concentrations of leucine and reduced growth in high concentrations of leucine. The difference at each concentration of leucine with and without BSA was highly significant (Ordinary one-way ANOVA, ****, p<0.0001). (F) Protein measurements taken at the end of the experiments shown in panel E, obtained from lysates of cells at 48 hrs. The difference with and without BSA was significant in some but not all concentrations of leucine (Student’s T-test of pairs at each leucine concentration; **, p<0.01, *, p<0.05, ns: not significant, n=9 wells per condition, 3 x 3 biological replicates).

To determine the range of leucine concentrations that support growth, we measured growth in R(0)/CSF1 supplemented with different concentrations of leucine. Incucyte growth curves showed limited growth at 1 and 3 mM leucine, intermediate levels of growth at 10 mM leucine, and maximal growth at leucine concentrations of 30 mM and higher (Fig. 2D). Robust growth at such low concentrations of leucine suggests that the volume of extracellular medium ingested by pinocytosis may be too small to deliver sufficient leucine for all of the protein synthesis in a growing macrophage.

Cell growth in R(BSA)/CSF1 indicated that hydrolysis of BSA in lysosomes can provide leucine for growth (Fig. 2A-C). To examine the contribution of added BSA to growth at low leucine concentrations, we measured macrophage growth in different concentrations of leucine, with and without 10 mg_*_ml^-1^ BSA. BSA increased growth in media with low concentrations of leucine but unexpectedly decreased growth in media with high concentrations of leucine (Fig. 2E). Direct measurements of protein content per well supported this conclusion (Fig. 2F). The rates of growth in BSA and leucine were consistent across the several experimental regimes (Suppl. Fig. 2B) and indicated that the effect of BSA on the rate of macrophage growth is dominant over the effect of free leucine on growth rates.

### Pinocytosis does not deliver sufficient leucine to support leucine-dependent growth

Cells in all growth media with CSF1 contained macropinosomes (Fig. 3A). To assess the effects of leucine on pinocytosis, LY accumulation was measured in R(0) and R(Leu), with and without CSF1 (Fig. 3B). Measurements of protein in wells containing different numbers of cells determined that macrophages were 118 pg protein_*_cell^-1^ in R(0)/CSF1, 127 pg protein_*_cell^-1^ in R(Leu)/CSF1 and 129 pg protein_*_cell^-1^ in R(BSA)/CSF1 (Suppl. Fig. 3A-C), which allowed calculation of rates of pinocytosis per 10^6^ cells in the various growth media. In the absence of CSF1, LY accumulation rates were low and unaffected by the presence or absence of leucine (Fig. 3C). CSF1 stimulated pinocytosis, and despite higher rates of cell growth in R(Leu)/CSF1 than in R(0)/CSF1 (Fig. 2C), LY accumulation was greater in the leucine-free medium. The linear accumulation rate in R(0)/CSF1 was greater than that in R(Leu)/CSF1 (Fig. 3C, D) or in complete medium (BMM, Fig 1G), indicating that cells drink more in medium lacking leucine. This result is consistent with an earlier study showing suppression of CSF1-stimulated pinocytosis by leucine (Mendel et al., 2022).

**Figure 3.**
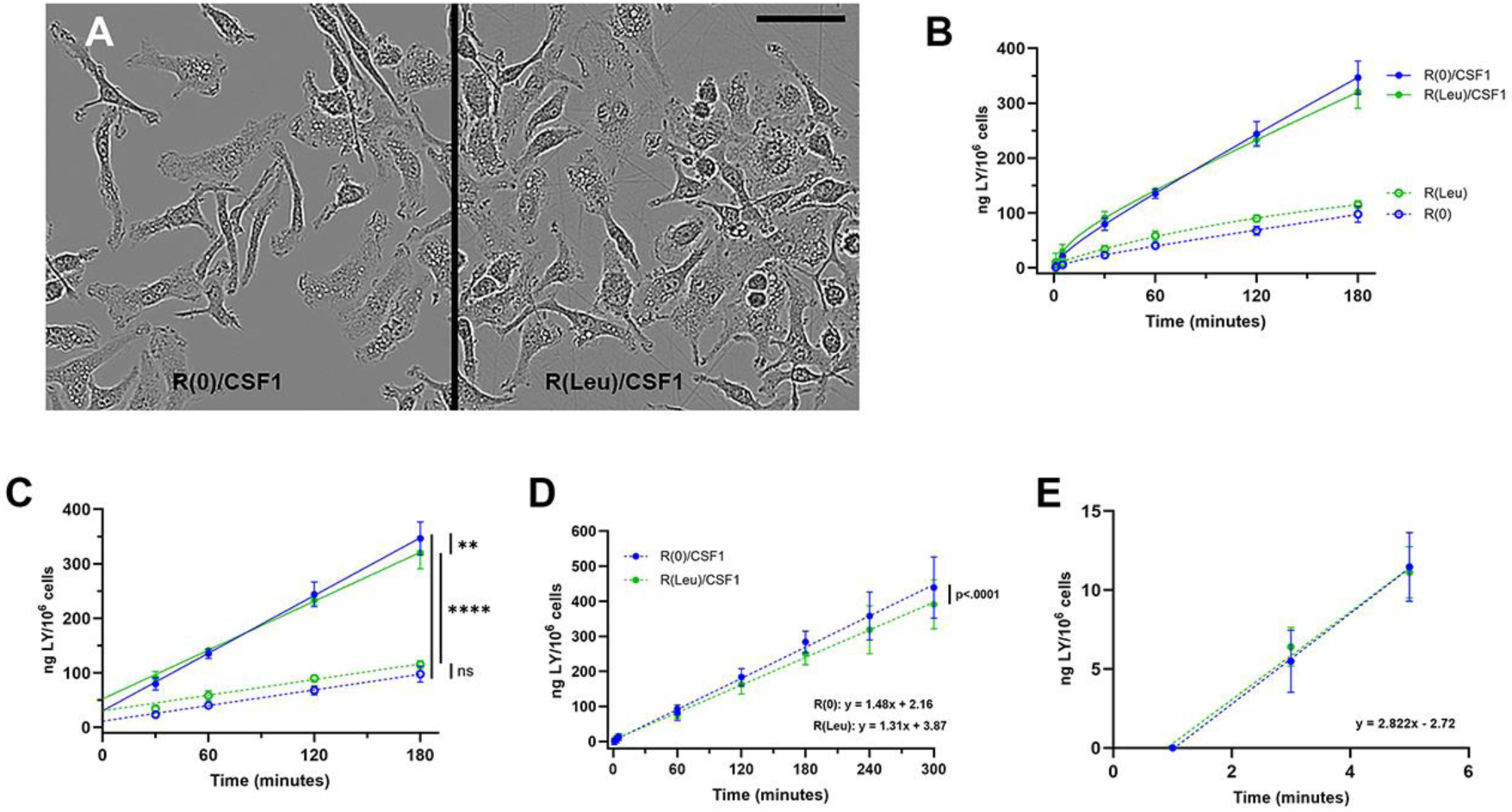
Absence of leucine increases rates of solute accumulation in growing macrophages. (A) Phase contrast images of macrophages in R(0)/CSF1 (left panel) and R(Leu)/CSF1 (800 mM leucine, right panel), obtained from the Incucyte at 30 hrs. Note phase-bright pinosomes and dividing cells in R(Leu)/CSF1. Scale bar: 50 mm. (B) Accumulation of 500 mg_*_ml^-1^ LY in R(0) (blue), +/- CSF1, +/- 800 mM Leu (green). (C) Rates of accumulation for the data in panel B (30-180 min). R(0): 0.490 ng LY_*_10^6^ cells^-1^ min^-1^; b=9.6, R^2^=0.92 (= 29.3 ng LY 10^6^ cells hr^-1^, = 58.7 fl fluid cell^-1^ hr^-1^); R(Leu): 0.532 ng LY_*_10^6^ cells^-1^ min^-1^; b=22.4, R^2^=0.94 (= 31.9 ng LY_*_10^6^ cells ^-1^ hr^-1^, = 63.8 fl fluid_*_cell^-1^ hr^-1^); R(0)/CSF1: 1.78 ng LY_*_10^6^ cells ^-1^ min^-1^; b=27.7, R^2^=0.97 (= 107 ng LY_*_10^6^ cells ^-1^ hr^-1^, = 214 fl fluid_*_cell^-1^ hr^-1^); R(Leu)/CSF1: 1.53 ng LY_*_10^6^ cells ^-1^ min^-1^; b=47.4, R^2^=0.97 (= 92 ng LY_*_10^6^ cells^-1^ hr^-1^, = 183 fl fluid cell^-1^ hr^-1^). One-way ANOVA of slopes: ns: not significant; **, p<0.01; ****, p<0.0001. (D) Rates of accumulation of 500 mg_*_ml^-1^ LY (1-300 min): R(Leu)/CSF1: 1.31 ng LY_*_10^6^ cells^-1^ min^-1^; b=0.07, R^2^=0.89 (= 79 ng LY 10^6^ cells^-1^ hr^-1^, = 157 fl fluid cell^-1^ hr^-1^), R(0)/CSF1: 1.48 ng LY_*_10^6^ cells^-1^ min^-1^ ; b=0.06, R^2^=0.89 (= 89 ng LY_*_10^6^ cells^-1^ hr^-1^, = 178 fl fluid_*_cell^-1^ hr^-1^). One-way ANOVA of slopes. (E) Rates of influx of 500 mg_*_ml^-1^ LY (accumulation at 1-5 min) were not significantly different for R(0)/CSF1 and R(Leu)/CSF1 (One-way ANOVA of slopes). The combined rates, obtained by subtracting the 1-minute values from the 3-and 5-minute values, were 2.82 ng LY_*_10^6^ cells^-1^ min^-1^, R^2^=0.90 (= 169 ng LY 10^6^ cells^-1^ hr^-1^, = 338 fl fluid cell^-1^ hr^-1^).

The rate of LY accumulation in R(Leu)/CSF1 was used to estimate leucine ingestion by pinocytosis. Accumulation of LY by cells in R(Leu)/CSF1 was 1.31 ng_*_10^6^ cells^-1^_*_min^-1^ (Fig. 3D). Influx in either R(0)/CSF1 or R(Leu)/CSF1 was 2.82 ng_*_10^6^ cells^-1^ min^-1^ (Fig. 3E), which translates to 338 fl of extracellular fluid ingested per cell per hour. The concentration of leucine in R(Leu)/CSF1 was 800 mM or 105 mg_*_ml^-1^. Assuming that ingestion of LY is proportional to the ingestion of leucine by fluid-phase pinocytosis, then macrophages in R(Leu)/CSF1 would have internalized 35.4 ng of leucine per 10^6^ cells per hour (Influx: 338 nl_*_10^6^ cells^-1^ hr^-1^ x 0.105 ng_*_nl^-1^ leucine). If all that leucine were used to make protein and if 5-10% of cellular protein is comprised of leucine, then pinocytosis of the 800 mM leucine in R(Leu)/CSF1 would allow a maximal synthesis of 354-708 ng protein per 10^6^ cells per hour, or up to 5574 cells_*_10^6^ cells^-1^ hr^-1^ (at 127 pg protein_*_cell^-1^). That would be insufficient to explain leucine-dependent growth in R(Leu)/CSF1 (20,300 cells_*_10^6^ cells^-1^ hr^-1^, Fig. 2C). The inadequacy of pinocytosis to deliver sufficient leucine to support growth was more pronounced in R(0)/CSF1 + 30 mM Leu (Fig. 2D), in which the 8-hr growth rates, normalized to the 30-hr time point, produced 22,800 cells_*_10^6^ cells^-1^ hr^-1^ but pinocytosis ingested only enough leucine to make protein for 105-210 cells 10^6^ cells^-1^ hr^-1^. Therefore, leucine-dependent growth cannot be supported solely by pinocytosis of leucine.

### Growing macrophages maintain high rates of solute efflux from lysosomes

A fraction of solutes internalized by pinocytosis are recycled to the medium. In studies of macrophages preloaded by pinocytosis with the fluid-phase solute probes ^14^C-sucrose and LY, two rates of efflux were observed: a rapid efflux (t_1/2_ = 5 min), which indicated recycling from early compartments, likely pinosomes, and a slower efflux (t_1/2_ = 180 min), that indicated recycling from lysosomes (Besterman et al., 1981; Swanson et al., 1985). To measure efflux during growth, macrophages were allowed to ingest LY at different concentrations in BMM for 2 hrs, then plates were washed free of LY and incubated 0, 1, 2 or 3 hrs in BMM. At the end of the chase periods, dishes were washed again, cells were lysed and the fluorescence of the remaining cell-associated LY was measured. The accumulation of LY over 2 hrs was proportional to the concentration of LY in BMM (0 min chase, Fig. 4A, Suppl. Fig. 4), confirming its utility as a bone fide probe of solute flux. The decrease in cell-associated LY, measured at 1, 2 and 3 hrs of chase, indicated a single exponential rate of dye efflux from lysosomes (Fig. 4B). However, plotting efflux as 1 – log^10^ fraction of the time-0 fluorescence remaining after 1, 2, and 3 hrs chase showed that rates of efflux were lower when cells were loaded with higher concentrations of LY (Fig. 4C). This suggests that the accumulation of LY in lysosomes interfered with its own recycling. A plot of the slopes in Fig. 4C vs. concentration of LY obtained a line with a y-intercept at -0.132 (antilog: 0.74) (Fig. 4D). This indicates that at vanishingly low concentrations of LY, macrophages in BMM lost 26% of their lysosomal LY per hour.

**Figure 4.**
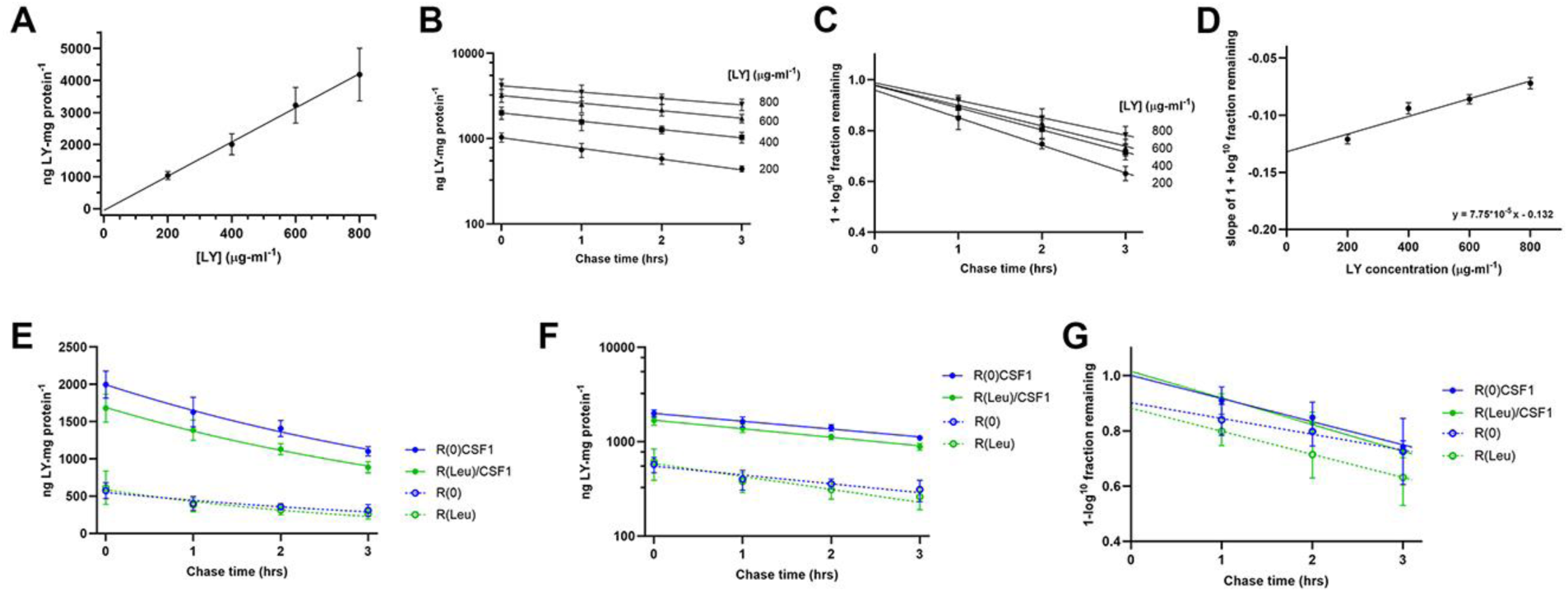
A large percentage of LY internalized by pinocytosis recycles from macrophages. (A) 2-hour accumulation of LY is proportional to the concentration of LY in BMM. Cells were incubated for 2 hrs in the indicated concentrations of LY before washing and lysis for determination of LY fluorescence and protein per well. These data are the 0-minute chase data for the efflux experiment (panels B-D). (B) Macrophages were labeled by pinocytosis of LY for 2 hr at the indicated concentrations in BMM, then were washed and chased for the indicated times in unlabeled BMM before washing and lysis of cells to measure retained LY. Rates of efflux from the cells indicate a first-order exponential decrease in cell-associated LY. (C) Fractional rates of LY loss from macrophages at different concentrations of LY. Cell-associated LY (ng LY_*_mg protein^-1^) was plotted as 1 – log^10^ of the fraction of LY remaining in the cells, relative to the 0-minute chase values for each concentration of LY. The slopes determined from linear regression analysis of 1, 2 and 3 hrs of chase in unlabeled medium indicate the rates of exponential loss of LY from the slowly emptying compartments (lysosomes). The y-intercept of those slopes indicates the relative sizes of the rapidly emptying compartments (pinosomes or endosomes). (D) Rates of slow efflux at different concentrations of LY. Slopes of 1 – log^10^ LY remaining at different chase times indicate slower rates of efflux from cells loaded with higher concentrations of LY. The y-intercept of the line described by those slopes, -0.132, indicates the rate of efflux at vanishingly small concentrations of LY. The antilog of that intercept is 26 % loss of LY per hr. (E, F, G) Effects of leucine and CSF1 on rates of LY efflux from pre-loaded lysosomes. (E) Cells growing in R(0) or R(Leu), with or without CSF1, were labeled by pinocytosis of 500 mg_*_ml^-1^ LY for 2 hrs, then were washed and returned to unlabeled media for 0, 1, 2, or 3 hrs before washing and lysis to measure LY retained in the cells. (F) Loss from the cells was exponential. (G) Linear regression of 1 – log^10^ fraction of 0-time values remaining in cells showed similar rates of slow efflux but significantly different y-intercepts with and without CSF1. Slopes for efflux in R(0)/CSF1 and R(Leu)/CSF1 showed y-intercepts of 1.0, whereas slopes for efflux in R(0) and R(Leu) had y-intercepts significantly less than 1.0, indicating the presence of a relatively large, rapidly emptying compartment.

To measure the effects of CSF1 and leucine on solute efflux, macrophages were allowed to pinocytose 500 mg_*_ml^-1^ LY for 2 hours in R(0) or R(Leu), +/- CSF1, then chased in similar media without LY for 0 to 3 hrs. Cells in leucine-containing media showed small but statistically insignificant increases in rates of slow efflux (Fig. 4E-G). The presence of CSF1 did not alter efflux rates, but decreased the size of the rapid efflux compartments, as indicated by the lower y-intercept of the efflux plots for cells without CSF1 (Fig. 4G). This indicates that CSF1 increases the percentage of internalized fluid solutes that are delivered into the slowly emptying lysosomes.

### Growth of macrophages requires macropinocytosis

To test the hypothesis that pinocytosis is required for cell growth, we measured the effects of pinocytosis inhibitors on macrophage proliferation in BMM. After testing many candidate inhibitors, we identified several that reliably inhibited pinocytosis of LY during a 5-hr incubation with macrophages (Fig. 5A). The most effective inhibitor was a combination of Jasplakinolide, which stabilizes actin filaments (Bubb et al., 1994), and para-amino-blebbistatin, which inhibits myosin II-based contractility (Straight et al., 2003; Varkuti et al., 2016). Consistent with previous studies, J/B reduced LY accumulation significantly (Lou et al., 2014; Yoshida et al., 2015). EIPA, a commonly used inhibitor of macropinocytosis, in our hands did not inhibit macropinocytosis significantly after the first hour. More significant inhibition of pinocytosis was obtained with SAR405, an inhibitor of type 3 PI-3-kinase (PI3K-3) (Ronan et al., 2014), and by bafilomycin A1, an inhibitor of the vacuolar ATPase (Bowman et al., 1988) (Fig. 5A). Because mTORC1 has been implicated in macropinosome-mediated degradation of protein in lysosomes (Palm et al., 2015), we also measured the effects of the mTOR inhibitors rapamycin and Torin-1 on pinocytosis. Rapamycin and Torin-1 stimulated LY accumulation (Fig. 5A).

**Figure 5.**
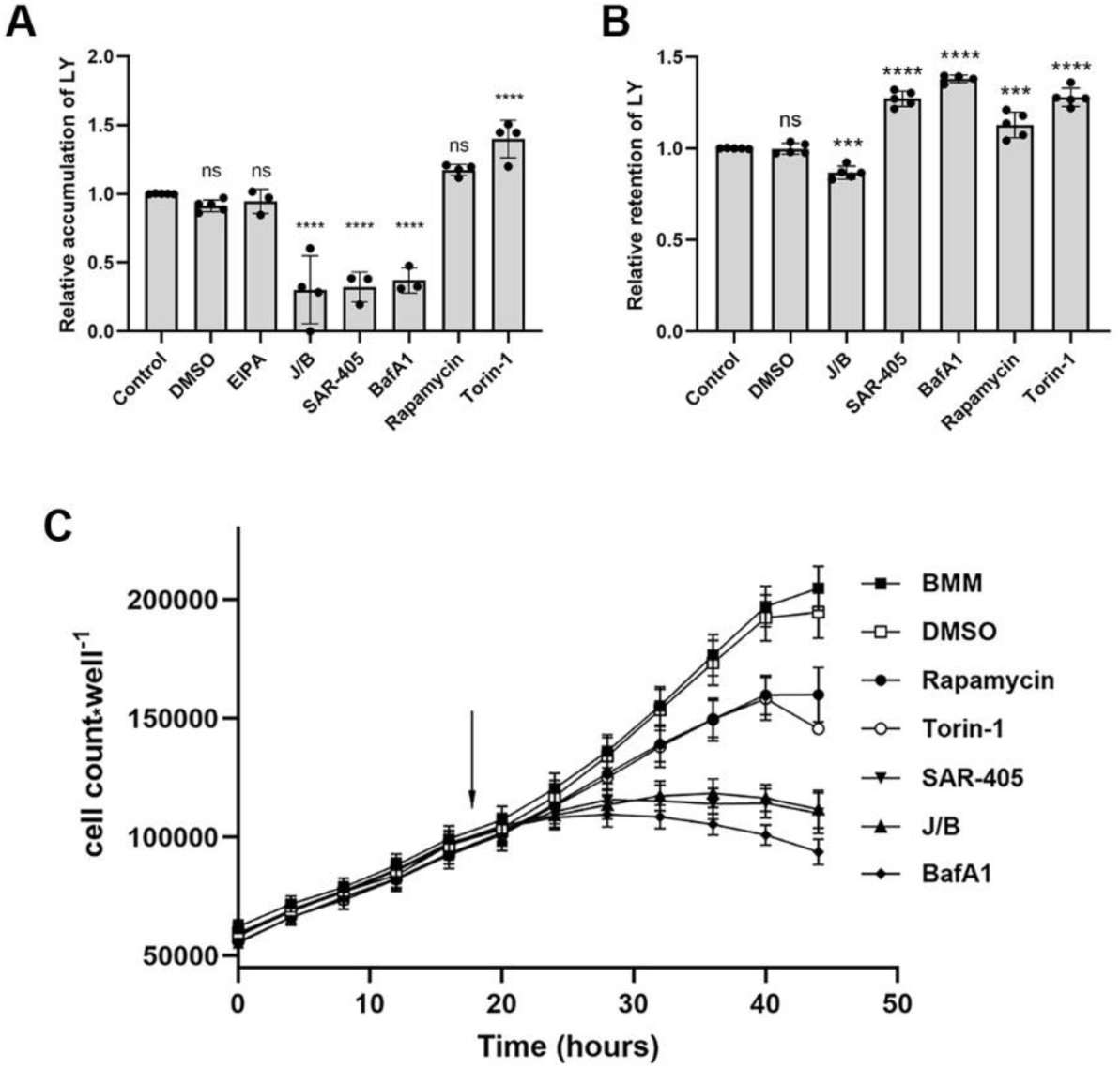
Inhibitors of pinocytosis and mTOR alter rates of LY accumulation and efflux. (A) Effects of inhibitors on LY accumulation by cells growing in BMM, measured after 5 hrs in 500 mg_*_ml^-1^ LY. Ordinary one-way ANOVA compared experimental to control values; ns, not significant; ****, p<0.0001. (B) Effects of inhibitors on LY efflux from lysosomes. Cells were pulsed 2 hr in BMM containing 500 mg_*_ml^-1^ LY, then were washed and chased 3 hr in BMM containing the indicated inhibitors. Dishes were then washed and lysed to measure remaining cell-associated LY. Higher values for retention indicate inhibition of recycling. J/B decreased LY retention and SAR-405, bafilomycin A1, Torin-1 and rapamycin increased retention. One-way ANOVA compared to control values; ns, not significant; ***, p<0.001, ****, p<0.0001. (C) Inhibitors were added to cells growing in BMM at 18 hr after the beginning of cell culture (arrow). Cell proliferation in BMM was inhibited by J/B, SAR-405, bafilomycin A1, rapamycin and Torin-1.

To measure effects of inhibitors on LY recycling from lysosomes, macrophages were labeled by incubation for 120 min in BMM containing LY, the plates were washed free of LY and cells were either lysed to measure LY content without a chase period or returned to warm BMM, +/- inhibitors, for 180 min before washing and lysis to measure LY remaining in cells. Relative to controls or to DMSO (vehicle)-treated cells, SAR405, bafilomycin A1, rapamycin and Torin-1 inhibited recycling (ie., increased retention of internalized LY). In contrast, J/B increased recycling (Fig. 5B).

The inhibitors of pinocytosis and mTOR altered cell morphology and inhibited macrophage growth. Macrophages growing in the Incucyte chamber were monitored for morphology and growth. After 18 hrs of growth in BMM, medium in wells was changed to BMM with or without inhibitors or vehicle, then monitored hourly thereafter (Fig. 5C). Images of cells after 3 hrs showed abundant macropinosomes in BMM, DMSO, rapamycin and Torin-1, extensive vacuolation in SAR405, and few or no pinosomes in J/B or bafilomycin A1 (Suppl. Fig. 5). Growth curves showed immediate inhibition of cell proliferation in J/B, SAR405 and bafilomycin A1 and a lesser inhibition of proliferation in rapamycin and Torin-1 (Fig 5C). Thus, pinocytosis is necessary for macrophage growth.

### Products of protein hydrolysis in lysosomes recycle from the cell

Rates of recycling from lysosomes vary inversely with solute size. Small molecules move between endocytic compartments and recycle from lysosomes more efficiently than do large solute molecules (Berthiaume et al., 1995). This suggests that peptides and amino acids released by hydrolysis of proteins in lysosomes may be excreted by growing macrophages. To test the hypothesis that small molecules generated by proteolysis in lysosomes are released from the cell, we measured the excretion of fluorescent molecules released by protein degradation in lysosomes. DQ-Red BSA is BSA covalently labeled with Bodipy TR-X to levels that quench its fluorescence (Vazquez and Colombo, 2009). Pinocytosis of DQ-Red BSA delivers it into lysosomes where it is degraded by lysosomal acid hydrolases, generating free, dequenched Bodipy TR-X. The fluorescent product colocalizes with LY delivered into lysosomes by pinocytosis (Suppl. Fig. 6A).

To measure recycling of small fluorophores generated by hydrolysis in lysosomes, macrophages were incubated 2 hours in medium containing DQ-Red BSA, LY, and Texas Red-labeled dextran, average molecular weight 10,000 (TRDx10) or 70,000 (TRDx70). Dishes were washed free of fluorophores and incubated in 0.5 ml Ringers buffer with BSA and CSF1 (R/B/C), for 0 or 180 minutes. Chase medium was then removed to measure fluorescence and cells on dishes were lysed to measure cell-associated fluorescence. Total fluorescence of LY, TRDx10 and TRDx70 (fluorophore in medium + fluorophore in cells) was the same at both chase times (Fig. 6A), confirming previous studies showing that LY fluorescence is not diminished inside the macrophages (Swanson et al., 1985). In contrast, the total fluorescence of DQ-Red BSA increased during the chase period due to the dequenching of the Bodipy TR-X upon its release by protein hydrolysis in the lysosomes (Fig. 6A). Moreover, the increase in total fluorescence of DQ-Red BSA was inhibited by adding 10 mM NH_4_Cl to the chase medium, which presumably increased lysosomal pH and inhibited hydrolysis and dequenching of DQ-Red BSA in lysosomes. For all probes, the fraction of total fluorescence in the medium increased during the chase period, indicative of recycling (Suppl. Fig. 6B). The fraction of total fluorescence recycled to the medium decreased with probe size. The fraction of LY recycled during the chase period was greater than the fraction of TRDx10 or TRDx70, which themselves recycled inversely with probe size (Fig. 6B). The Bodipy TR-X released by proteolysis of DQ-Red BSA recycled as efficiently as LY, indicating that the products of lysosomal proteolysis are released from the cells at rates inversely proportional to molecular size. These unexpectedly high rates of efflux suggest that macrophages either secrete small nutrient molecules generated by lysosomal hydrolysis, such as amino acids, or actively retrieve essential nutrients on the route out of the cells.

**Figure 6.**
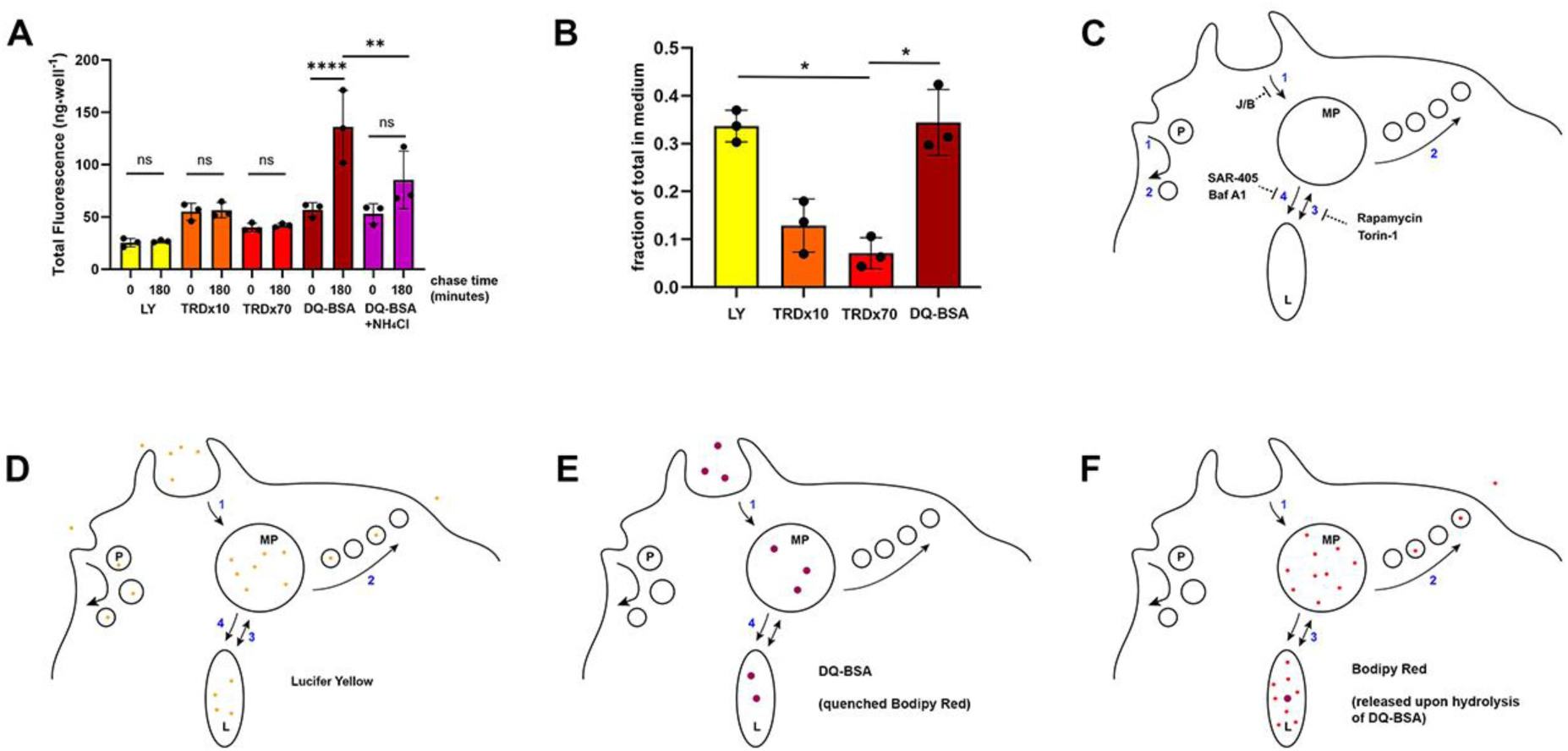
Small molecules generated by protein degradation in lysosomes recycle from the cell. (A, B) Efflux of LY, TRDx10, TRDx70 and DQ-BSA (Bodipy Red). (A) Macrophages were incubated with the indicated fluorophores for 2 hr in BMM, then were washed and the medium was replaced with Ringers Buffer with BSA and CSF1 for 0 or 180 min. Chase medium was removed and cells were lysed to measure cell-associated fluorescence. Total fluorescence in the chase medium and cell-associated fluorescence were calculated for each well. Total fluorescence of LY, TRDx10 and TRDx70 were constant, confirming that the probes were not altered or degraded in the cells. The fluorescence of DQ-BSA increased significantly due to dequenching of the fluorophore Bodipy Red upon its release from the BSA by lysosomal hydrolysis. That increase was diminished by inclusion of 10 mM NH_4_Cl in the chase medium. One way ANOVA of the indicated pairs: ns: not significant, **, p<0.01; ****, p<0.0001. (B) The fraction of internalized probe that recycled to the medium was inversely proportional to the probe size. Recycling efficiency of Bodipy Red (from DQ-BSA) resembled the efficiency of the small probe LY. (C) Model for pinocytic solute influx and retrograde flow during cell growth. In the absence of CSF1, most pinocytosis is via small pinosomes (P) which recycle fluid and send relatively small quantities of ingested solutes to lysosomes. CSF1 stimulates macropinosome formation (MP) by macropinocytosis (1), which can be inhibited by J/B. Membrane and a small fraction of internalized solutes recycle from macropinosomes in small vesicles (fast recycling, 2). Macropinosomes interact with lysosomes either by pyranhalysisis, with transient small bridging connections between the organelles (3) or by complete merger of macropinosome content and membrane into lysosomes (4). We propose that the former is inhibited by rapamycin and Torin-1 and the latter is inhibited by SAR-405 and Bafilomycin A1. (D) Lucifer yellow reports movement through all these pathways. (E) Being larger than LY, DQ-BSA enters primarily in macropinosomes and is restricted by its size-limited traffic itinerary to the more direct fusion with lysosomes. (F) Upon degradation of DQ-BSA in lysosomes and release of Bodipy TR-X into the lumen, the smaller fluorophore recycles efficiently from the cell, moving retrograde via size-selective interactions with macropinosomes.

## Discussion

Although macropinocytosis has long been recognized as an acute response to growth factor stimulation, no prior studies have analyzed the contributions of growth factor-stimulated macropinocytosis to growth. Here we show that macrophage growth is accompanied by high rates of pinocytic solute influx and efflux, driven by CSF1 signaling and regulated by leucine. The high rates of solute influx and accumulation are not sufficient to internalize all the leucine required for growth, but pinocytosis is nonetheless necessary for growth and proliferation.

### Solute influx and accumulation are regulated by CSF1 and leucine

CSF1 is the principal regulator of pinocytosis in macrophages. It stimulates macropinocytosis and decreases the percentage of internalized LY that recycles from the cell, leading to dramatic increases in the rates of solute accumulation (Racoosin and Swanson, 1989). The present study indicates that stimulation with CSF1 does not alter the rate of slow recycling, but rather increases the percentage of internalized LY that recycles quickly from the cell. The y-intercept of the plot of 1-log^10^ fraction remaining vs. chase time (Fig. 4G) shows that in CSF1-containing media all of the LY in the cells recycles at the slow rate (y-intercept = 1.00), whereas in CSF1-deficient media only 90% of the internalized LY recycles at the slow rate (y-intercept = 0.90). Pinocytosis in CSF1 takes in a large volume of fluid and sends it all to the lysosomes.

Although less dramatic than the effects of CSF1, leucine also regulates solute flux. Rates of LY influx were nearly identical in R(0)/CSF1 and R(Leu)/CSF1 (Fig. 3E), but net accumulation was greater in R(0)/CSF1 (Fig. 3C, D), which indicates that recycling was inhibited by the lack of leucine. Leucine and other suppressors of pinocytosis reduce macropinosome size, in part by promoting the loss of CSF1 receptors (Mendel et al., 2022). In the absence of leucine, accumulation must increase by a small difference in rates of influx or recycling. In their analysis of the effects of EAA on pinocytosis in rat alveolar macrophages, Besterman et al. found that the presence or absence of EAA did not affect rates of slow or fast solute recycling of ^14^C-sucrose; however, the absence of EAA increased the size of the slow recycling compartment (Besterman et al., 1983). This indicates that more of what entered the cell by pinocytosis moved from the rapidly recycling compartment to the slowly recycling compartment, resulting in a net increase in accumulation. A similar shift in the proportions of internalized LY moving between pinosomes and lysosomes may explain the increased LY accumulation in the absence of leucine.

### CSF1-stimulated pinocytosis does not internalize enough fluid to support leucine-dependent growth

Extracellular leucine must be imported across membranes into cytoplasm for it to be available for protein synthesis. That import could be across plasma membrane, lysosome membrane or macropinosome membrane. In macrophages growing in 30 mM leucine, the cells’ capacity for pinocytosis could not internalize sufficient free leucine to support protein synthesis. The amino acid probably enters cytoplasm mostly by leucine transport proteins in plasma membranes. The fact that inclusion of BSA in media overrides effects of leucine concentrations on growth rates (Fig. 2E, F) suggests that cells can monitor and regulate their sources of leucine between transporters in plasma membrane or pinosomes and hydrolysis of internalized proteins in lysosomes.

### Efflux of small solutes indicates a major pathway of solute flow in growing macrophages

The dynamics of pinocytosis in growing macrophages suggest a model for the regulation of solute and nutrient flux (Fig. 6C). Membrane-impermeant, fluid-phase solutes can enter cells by small pinosomes (P), which form constitutively, or by larger macropinosomes (MP), which assemble in response to CSF1 (Fig. 6C, step 1). Macropinosomes form primarily by activities of the actin cytoskeleton, which can be inhibited by J/B (Lou et al., 2014; Yoshida et al., 2015). Small solutes recycle and move more easily between endocytic compartments than do large solutes (Berthiaume et al., 1995), and we propose that after macropinosomes form they shrink and recycle small solutes through small vesicles or tubular extensions (Freeman et al., 2020; Swanson, 1989) (Fig. 6C, step 2). Further, macropinosomes interact with lysosomes (L) by pyranhalysis (Willingham and Yamada, 1978), exchanging small molecules between the two compartments more readily than large molecules (Berthiaume et al., 1995; Yoshida et al., 2015) (Fig. 6C, step 3). Slow recycling of small molecules from lysosomes likely occurs via these transient interactions with macropinosomes, allowing small solutes to leave the cell via steps 2 and 3. Sometime later in its maturation, the macropinosome merges into the lysosomes completely, delivering the remaining mixture of large and small molecules (Fig. 6C, step 4).

Solute size-selective exchange (step 3) followed by full merger of pinosome contents into lysosomes (step 4) would allow a net accumulation of large molecules, like proteins, and a net retrograde flow of small solutes, like amino acids derived from protein hydrolysis. It is also possible that slow efflux occurs by direct communication of lysosomes with the plasma membrane (Andrews, 2000; Besterman et al., 1981). However, the fact that bafilomycin A1 and SAR405 inhibit both accumulation and efflux suggests that one interaction regulates flow in both directions. Thus, we propose that the component activities of macropinocytosis are (1) influx, (2) recycling from pinosomes, (3) pyranhalysis and (4) merger with lysosomes.

The small probe LY reports the movements of fluid-phase solutes through this system of organelles (Fig. 6D). It labels small pinosomes and macropinosomes, recycling compartments and lysosomes and it effectively measures influx and accumulation (Swanson et al., 1985). LY reports high rates of efflux from lysosomes, higher rates than are observed with larger probes (Berthiaume et al., 1995)(Fig. 6B), which is consistent with pyranhalysis as a component of pinosome-lysosome interactions (step 3).

The dynamics of TRDx10, TRDx70 and DQ-BSA in macrophages indicate the size-dependence of movements into and between these organelles (Fig. 6E). Larger probes of pinocytosis such as these would move less readily into small vesicles and between macropinosomes and lysosomes. Size-selective retrograde flow of LY and TRDx10 from lysosomes into macropinosomes has been reported (Yoshida et al., 2015). A macropinosome maturation step that includes full merger into lysosomes (step 4), like phagosome-lysosome fusion, would permit robust accumulation of proteins and larger molecules in lysosomes. Such movements have also been observed (Yoshida et al., 2015). Degradation of DQ-BSA and the attendant release of Bodipy Red would allow increased recycling of the smaller dye out of the cell (Fig. 6F).

The inhibitors of macropinocytosis and mTOR have different effects on pinocytosis and recycling, therefore they likely act at different stages of the process. Pinocytosis inhibitors could be limiting either a primary route of some nutrients into cells, the traffic of receptors or amino acid transporters from intracellular compartments to the cell surface, or some combination of these effects. J/B inhibited pinocytosis and increased LY recycling from lysosomes, which suggests that the actin cytoskeleton is necessary for communication between pinosomes (compartment 1) and lysosomes (compartment 2). Accordingly, J/B may cause more internalized LY to recycle from pinosomes than to move into lysosomes. Bafilomycin A1 elevates the pH of acidic organelles (Bowman et al., 1988) and inhibits endocytic membrane traffic (Mauvezin et al., 2015). SAR405 inhibits VPS34, the type III PI-3-kinase that regulates endosome-lysosome fusion, membrane recycling and autophagy (Backer, 2008). Because bafilomycin A1 and SAR405 inhibit LY accumulation and efflux, we suggest that they prevent the merger of pinosomes into lysosomes (step 4). In contrast, rapamycin and Torin-1 stimulate accumulation and inhibit recycling, which suggests that they do not prevent the merger of macropinosomes into lysosomes but rather the size-selective recycling of solutes (step 3). These speculations support the model of Figure 6C, but certainly more work will be needed to dissect the component activities behind solute dynamics.

At the concentrations used to measure pinocytosis and efflux, LY may interfere with the dynamics of solute recycling. Higher concentrations of LY reported slower rates of recycling than did low concentrations (Fig. 4C, D). The rate of slow efflux indicated for the absence of LY -the y-intercept of efflux rates vs. LY concentrations (Fig. 4D) -is 26% of the lysosomal LY every hour. That graph also indicates that the 500 mg_*_ml^-1^ LY used to measure solute flux should recycle LY-size molecules from lysosomes at 19% per hour.

The high rates of small molecule recycling indicate that lysosome recycling is a significant and underappreciated activity of macrophages and growing cells. Perhaps pinocytosis by macrophages of serum proteins, degradation and subsequent recycling provides EAA or other products of lysosomal hydrolysis that support the growth of nearby cells. Alternatively, small nutrients such as amino acids recycling from lysosomes into macropinosomes could be retrieved into cytoplasm on their way out of the cell by transport proteins in the membranes of macropinosomes. Transporters in macropinosome membranes could utilize gradients of H^+^, Na^+^ or Cl^-^ in the volume of ingested medium to facilitate nutrient import. The macropinosome could thus serve as a single port of nutrient entry into cytoplasm, importing both the nutrients engulfed into the pinosome and those that recycle from lysosomes.

What role, if any, does mTORC1 serve in macropinocytosis? mTORC1 can be activated by leucine in macropinosomes (Yoshida et al., 2015), which suggests that mTORC1 regulates macropinosome function. Active mTORC1 inhibits degradation of proteins ingested by macropinocytosis (Palm et al., 2015), in part by suppressing assembly and activity of the V-ATPase and the consequent hydrolysis of proteins in lysosomes (Ratto et al., 2022). An alternative explanation may be that mTORC1 inhibits macropinocytosis, either through inhibition of the V-ATPase (the target of Bafilomycin A1) or by stimulating solute recycling.

Macropinocytosis of extracellular proteins can support the growth of Ras-transformed cancer cells in nutrient-poor tumor environments (Commisso et al., 2013). The dynamics of solutes indicated by this study suggest that amino acids generated by lysosomal hydrolysis of proteins scavenged by cancer cells may be recycled to macropinosomes or secreted from the cells. The magnitude of recycling by small molecules from lysosomes suggests that the process may be important for normal physiology and possibly altered in methuosis, a condition in which macropinocytosis kills the cell (Maltese and Overmeyer, 2015; Ritter et al., 2021).

## Materials and Methods

### Bone Marrow-derived Macrophages

Bone marrow-derived macrophages were obtained from C57BL/6J mice (Jackson Labs, Bar Harbor, ME). Marrow was flushed from femurs and cultured for 6 days in bone marrow medium (BMM): RPMI medium supplemented with recombinant murine CSF1 (50 ng/ml; R&D Systems, #416-ML-500), 50 mM b-mercaptoethanol, 23.8 mM sodium bicarbonate, 20% heat-inactivated fetal bovine serum (Life Technologies Corp. #16000044) and 0.1% Penicillin/Streptomycin. On the third day of cell culture, supernatant containing floating macrophage colonies was transferred into 100 mm diameter plates to obtain synchronously developing, adherent macrophages. On the sixth day, media was replaced with cold PBS for 10-20 min at 4^0^ C followed by pipetting to resuspend cells. Cells were pelleted by centrifugation, resuspended in BMM, then plated into 24-well tissue culture dishes (Corning, USA), or into 35 mm glass-bottom microwell dishes (MatTek corporation, Ashland MA). Plated cells were cultured in BMM or other defined media, with or without CSF1 (50 ng/ml), for measurements of growth and pinocytosis. All animal-related procedures were approved by the University of Michigan Committee on Use and Care of Animals.

For live cell imaging of macrophages, MatTek dishes were maintained at 37°C in Ringer’s Buffer (RB: 155 mM NaCl, 5 mM KCl, 2 mM CaCl_2_, 1 mM MgCl_2_, 2 mM NaH_2_PO_4_, 10 mM glucose and 10 mM HEPES at pH 7.2) supplemented with 1 mg/ml BSA and 50 ng/ml CSF1 (R/B/C).

### Cell growth media and inhibitors

To study leucine-dependent growth, a growth medium lacking leucine was prepared (R(0)): to RPMI 1640 powder without arginine, leucine or lysine (US Biological Life Sciences, #R8999-03A) were added L-arginine-HCl (Sigma #A5606, 200 mg_*_l^-1^, 1.15 mM), L-lysine-HCl (Sigma, L5501, 40 mg_*_l^-1^, 0.22 mM), 10% heat-inactivated, dialyzed fetal bovine serum (dFBS, ThermoFisher Scientific. #A3382001), sodium bicarbonate (23.8 mM), b-mercaptoethanol (50 mM), and Penicillin/Streptomycin (0.1%). R(0) was then supplemented to obtain media containing 800 mM leucine (R(Leu)), 800 mM isoleucine (R(Iso)), 10 mg_*_ml^-1^ endotoxin-free bovine serum albumin (R(BSA), Sigma Aldrich #A7906) or 50 ng/ml CSF1 (Recombinant Mouse M-CSF, R&D Systems 416-ML-500)(R(0)/CSF1, R(Leu)/CSF1, R(Iso)/CSF1, R(BSA)/CSF1) by combining concentrated stock solutions of these ingredients prepared in R(0).

Inhibitors from concentrated stock solutions in DMSO were prepared in BMM or R(0) for measurements of growth and solute flux by pinocytosis, including EIPA (25 mM final, Cayman Chemical, #14406), jasplakinolide (1 mM final, Cayman Chemical #11705), para-amino blebbistatin (75 mM final, Cayman Chemical #22699), bafilomycin A1 (500 nM final, Tocris/Biotechne #3872), SAR405 (10 mM final, Selleckchem #S7682), rapamycin (100 nM final, Tocris/Biotechne #1292), and Torin-1 (1 mM final, Tocris/Biotechne #4247), with DMSO as vehicle controls.

### Measurement of cell growth

Cell growth was measured using an Incucyte SX1 (Sartorius), a system in which a microscope inside a 37°C CO_2_ incubator scans cell culture samples repeatedly over pre-determined intervals to obtain images of cell fields and, from them, to generate quantitative kinetic measurements of cell growth and proliferation. 24-well plates containing 8-10 x 10^4^ cells per well were cultured in different media according to the experiment. Nine different regions of each well were imaged every hour for 48 hours. Cell growth was measured using Incucyte software that determined cell numbers in each well of the plate (Cell-by-Cell, Sartorius). Rates of growth were uniform from 24-40 hrs of culture. To calculate growth rates, cell numbers per well, measured from 30-38 hr of culture in the Incucyte SX1, were normalized to the numbers at 30 hrs for each condition. Normalized cell growth was obtained by linear regression analysis of the normalized cell numbers per well vs. time. The slopes of those lines were reported as the number of cells produced per hour per 10^6^ cells at the 30-hr time point. Those rates were multiplied by the calculated protein per cell (see below) to obtain the rates of protein synthesis in each condition.

Growth was also measured as the increase in cellular protein, using the Pierce bicinchoninic acid (BCA) protein assay (Life Technologies Corp. #23225). Cells in 24-well dishes were lysed in 0.4 ml 0.1% Triton X-100 and immediately analyzed for protein content. This method was not as sensitive as the Normalized Cell Growth method described above.

### Measurement of protein per cell

Various numbers of cells were plated into wells of a 24-well dish and cultured overnight in BMM/CSF1, R(0)/CSF1, R(Leu)/CSF1 or R(BSA)/CSF1. Dishes were washed in cold PBS to remove medium and serum and then returned to the Incucyte chamber in RB. Nine frames per well were collected from each well and the cell numbers per well were calculated using the Cell-by-Cell image analysis program. RB was removed and cells were lysed for measurement of protein content per well (BCA assay). Linear regression analysis of cell number per well vs. mg protein per well allowed calculation of the pg protein per cell in each growth medium. Data were compiled from 3 biological replicates of the experiment.

### Measurement of pinocytosis

The movements of solutes into and through macrophages were measured using Lucifer yellow CH (LY, Invitrogen), as described previously (Swanson et al., 1985). Macrophages on day 6 of culture were plated at 1 x 10^5^ cells/well in 24-well dishes in BMM, R(0) or R(Leu), +/-CSF1, and were incubated overnight at 37 ^0^C. Cells were incubated in media containing 500 mg_*_ml^-1^ LY (0.35 ml/well) for 1-300 minutes at 37^0^C, such that all incubations ended simultaneously. LY-containing medium was shaken from the plates, then plates were washed by immersion into 1 liter of cold PBS with 1 mg/ml BSA and then into two sequential liters of ice-cold PBS. Dishes were drained and 0.6 ml of Lysis buffer (0.1% Triton X-100, 0.02% sodium azide) was added to each well to lyse cells. To measure the amount of LY that had accumulated inside the cells, 0.5 ml of lysate was combined with 0.4 ml of Lysis buffer with 1 mg/ml BSA (LB/BSA) and fluorescence of LY was measured in a fluorometer (Duetta, Horiba Scientific); exc. 430 +/- 5 nm, em. 540 +/- 5 nm. LY standard curves were prepared by dilution of the drink solution. 70 ml of lysate was removed to measure the mg protein per well, using the BCA assay. The amount of LY ingested per mg of protein was calculated. To determine the rate of pinocytosis per 10^6^ cells, the protein content of each well was converted to cell number per well using the calculated determinations of protein per cell (Figs 1D, Suppl. Fig. 3A, B, C).

### Efflux measurements

Recycling of LY from cells in BMM was measured by allowing cells plated into 12 wells of 4 24-well dishes to internalize 200 mg_*_ml^-1^, 400 mg_*_ml^-1^, 600 mg_*_ml^-1^ or 800 mg_*_ml^-1^ LY in BMM/CSF1 for 2 hrs (each dish contained all four concentrations of LY). Dishes were emptied and washed by immersion in 2 liters of cold PBS, then were either lysed (dish 1, for 0-minute chase values) or were returned to unlabeled BMM/CSF1 for 1, 2 or 3 hrs (dishes 2-4) before washing again in cold PBS and lysing in Lysis buffer to determine LY and protein content per well (ng LY_*_mg protein^-1^). Efflux of LY was measured as the decrease in cell-associated LY as a function of time.

Recycling of LY from macrophages in R(0), R(Leu), R(0)/CSF1 and R(Leu)/CSF1 was measured after first transferring 4 dishes with 12 wells of cells per dish to the 4 different media for 3 hrs. Cells were then incubated for 2 hrs in the corresponding growth media containing 500 mg_*_ml^-1^ LY. Dishes were washed to remove LY and either lysed (dish 1, chase time = 0 hrs) or returned to LY-free growth media for 1, 2 or 3 hrs (dishes 2-4) before washing again and lysing to measure retained LY and protein per well.

To measure effects of inhibitors on efflux, macrophages in two 24-well dishes were incubated 2 hrs in BMM containing 500 mg_*_ml^-1^ LY. Dishes were washed free of LY and medium, then either lysed (dish 1, 0-minute chase) or returned to BMM containing DMSO (vehicle control) or the inhibitors. After 3 hrs of chase, dish 2 was washed in cold PBS, cells were lysed, and remaining LY and protein per well were determined. Increased content of LY, relative to cells chased in BMM, indicated inhibition of LY efflux.

To measure efflux of different sizes of fluid-phase probe and of DQ-Red BSA, macrophages in 2 24-well dishes were incubated for 2 hrs in BMM/CSF1 containing 500 mg_*_ml^-1^ LY +/- TRDx10 (Thermo Fisher Scientific; 100 mg_*_ml^-1^), TRDx70 (Thermo Fisher Scientific; 100 mg_*_ml^-1^), or DQ-Red BSA (Thermo Fisher Scientific; 10 mg_*_ml^-1^). Dishes were washed in 2x 1 liter PBS, drained, then replenished with 0.5 ml R/B/C for 0 or 3 hrs at 37 C. Medium was collected from wells and combined with 0.5 ml LB/BSA. Cells in wells were lysed in 0.5 ml lysis buffer and combined with 0.5 ml LB/BSA. LY fluorescence was measured at exc. 430 +/- 5 nm, em. 540 +/- 5 nm. TR and Bodipy TR-X fluorescence were measured at exc. 588 +/- 5 nm, em. 616 +/- 5 nm. The LY and TR/BodipyTR-X signals were not detectable in the other fluorophore’s channels.

### Microscopy

Fluorescent cells were imaged in a Nikon Eclipse TE-300 inverted fluorescence microscope with a 60x NA 1.4, oil-immersion Plan Apo objective lens and a Lambda LS xenon arc lamp for epifluorescence illumination (Sutter Instruments). Fluorescence excitation and emission wavelengths were selected using a 69008 filter set (Chroma Technology) and a Lambda 10-2 filter wheel controller (Sutter Inst.) equipped with a shutter for epifluorescence illumination control. Images were recorded with PCO.Edge sCMOS camera (Mager Scientific).

For live cell imaging, cells plated onto glass bottom, 35 mm diameter dishes (MatTek Corp.) were preloaded by endocytosis of LY (500 mg_*_ml^-1^) and/or DQ-Red BSA (10 mg_*_ml^-1^) in BMM for 2 hrs, then were washed in cold PBS and incubated in RB/BSA/CSF1 at 37 C. Images were processed using Metamorph software (Molecular Devices).

### Statistical analysis

All measurements were obtained from triplicate measurements of 3-5 biological replicates. Quantitative data were analyzed first in Excel, then exported to Prism for statistical analysis and graphical presentation. Data were analyzed by simple linear regression or by one-way ANOVA. Statistical significance of differences are indicated in the figure legends.

## Supplemental materials

Supplementary figures show: pinocytosis measurements replotted as ng LY per mg protein (Suppl. Fig. 1), rates of cell growth when isoleucine is substituted for leucine (Suppl. Fig. 2A), compiled rates of growth in different media (Suppl. Fig. 2B), measurements of protein content per cell in R(0), R(Leu) and R(BSA) (Suppl. Fig. 3), LY concentration-dependence of accumulation (Suppl. Fig. 4), images from the Incucyte of cells in various inhibitors (Suppl. Fig. 5), fluorescence colocalization of LY and DQ-Red BSA (Suppl. Fig. 6A), and fluorescence of dyes recycled from lysosomes at 0- and 180-minutes chase times (Suppl. Fig. 6B).

## Acknowledgment

This work was supported by the University of Michigan Medical School and funds from the NIH (R35-GM131720 to J.A.S. and R15-GM139162 and P20-GM135008 to N.W.T). The authors are grateful for the suggestions of Drs. Adam Hoppe and Philip King.

---

**Supplementary Figure 1.**
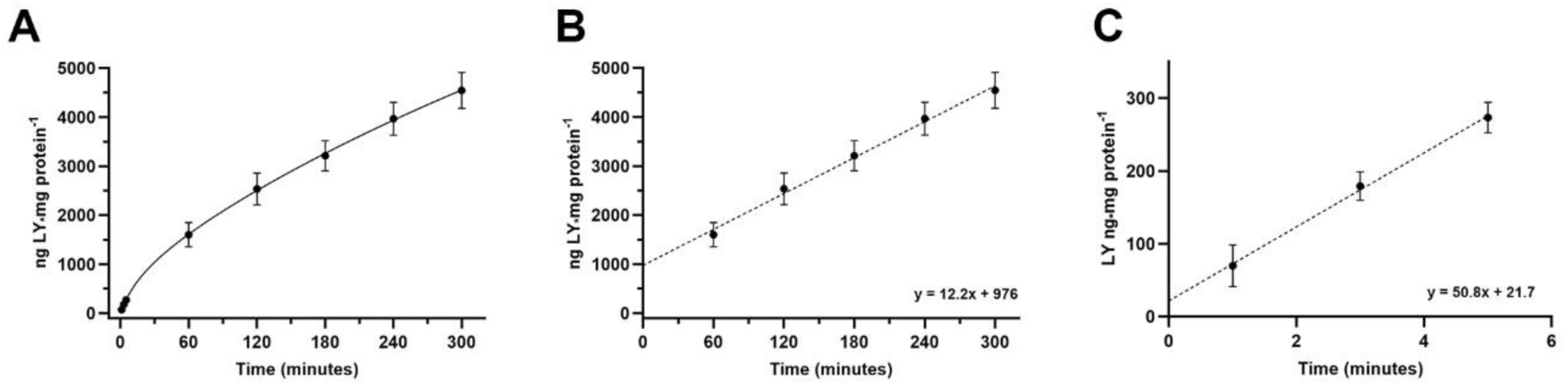
The data of Figure 1 B-D plotted as ng LY_*_mg protein^-1^, as in previous publications (Berthiaume et al., 1995; Swanson et al., 1985).

**Supplementary Figure 2.**
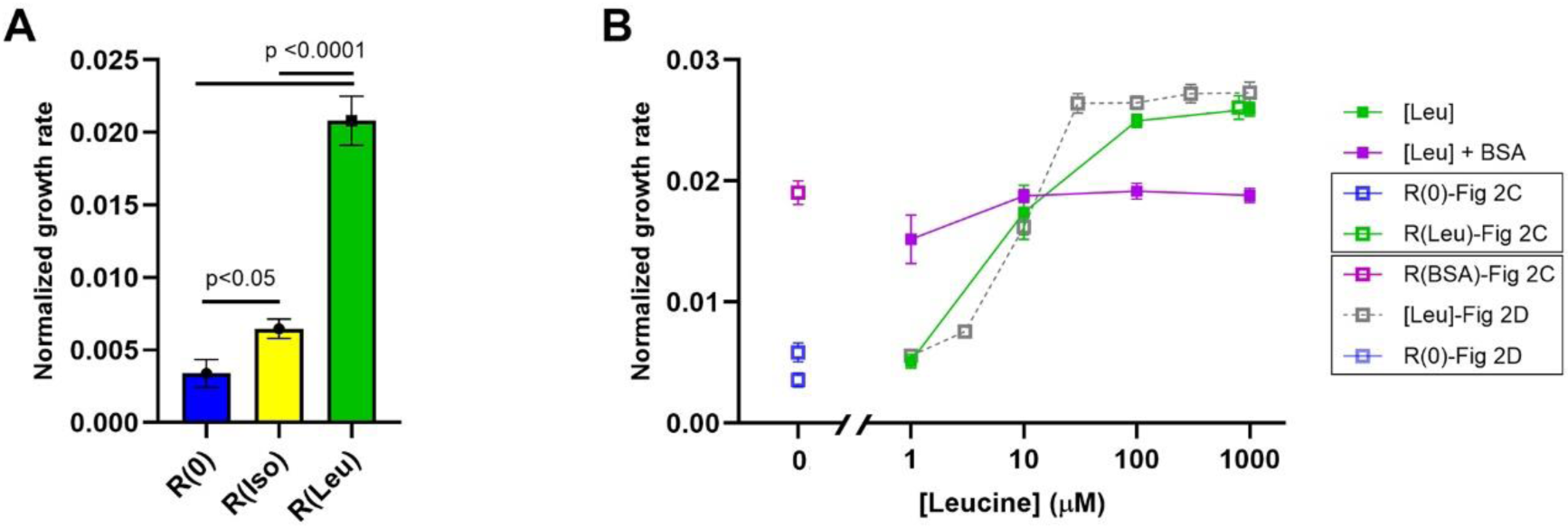
(A) Rates of cell growth in R(0)/CSF1, R(Iso)/CSF1, R(Leu)/CSF1, showing mean and SEM for rates of growth normalized to the cell counts for each condition at 30 hrs. One-way ANOVA, n = 81 per condition (9 time points from 30-38 hrs, 3 wells per time point, 3 biological replicates). (B) Compiled data from related experiments that measured rates of cell growth in various media, normalized to the 30-hour time point. Solid squares indicate data of Fig. 2E. Open squares indicate data of Fig. 2C and Fig 2D.

**Supplementary Figure 3.**
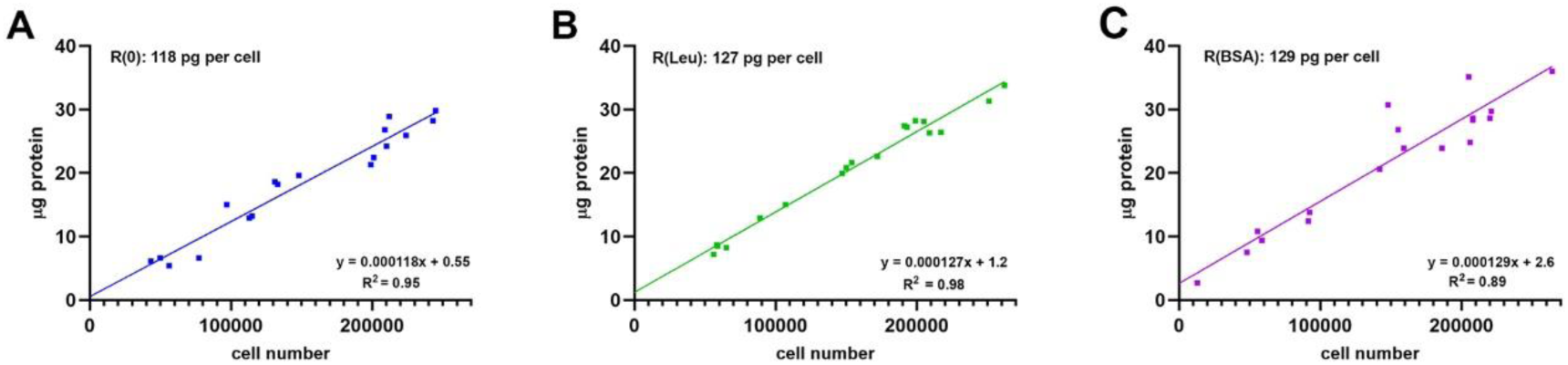
Plots of mg protein_*_well^-1^ vs. cell numbers obtained from the Incucyte allow determination of protein content per cell in the various growth media. (A) R(0), (B) R(Leu), (C) R(BSA).

**Supplementary Figure 4.**
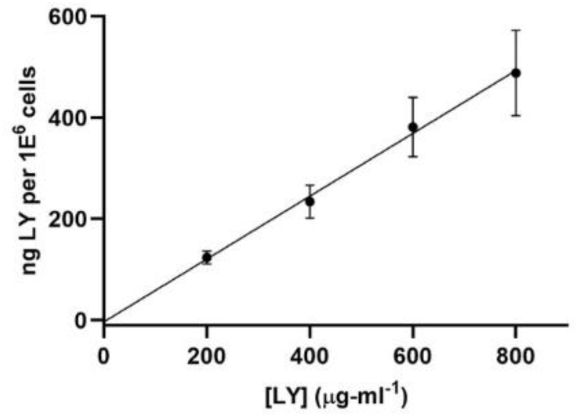
The data of Fig. 4A recalculated to show ng LY_*_10^6^ cells^-1^.

**Supplementary Figure 5.**
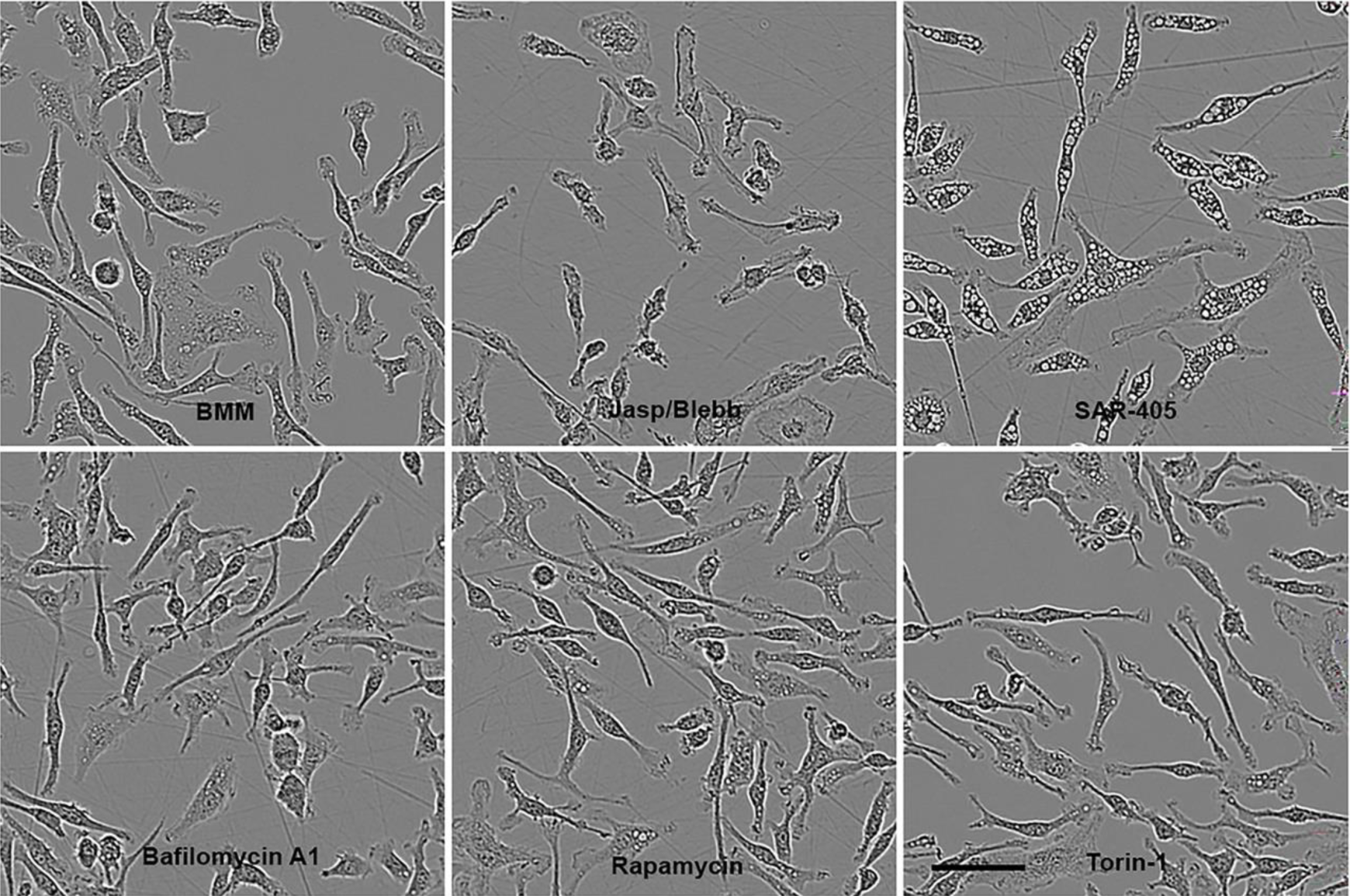
Effects of inhibitors on macrophage morphology obtaine from incucyte images taken 3 hr after addition of BMM (control), J/B, SAR-405, Bafilomycin A1, Rapamycin and Torin-1. Scale bar: 50 mm.

**Supplementary Figure 6.**
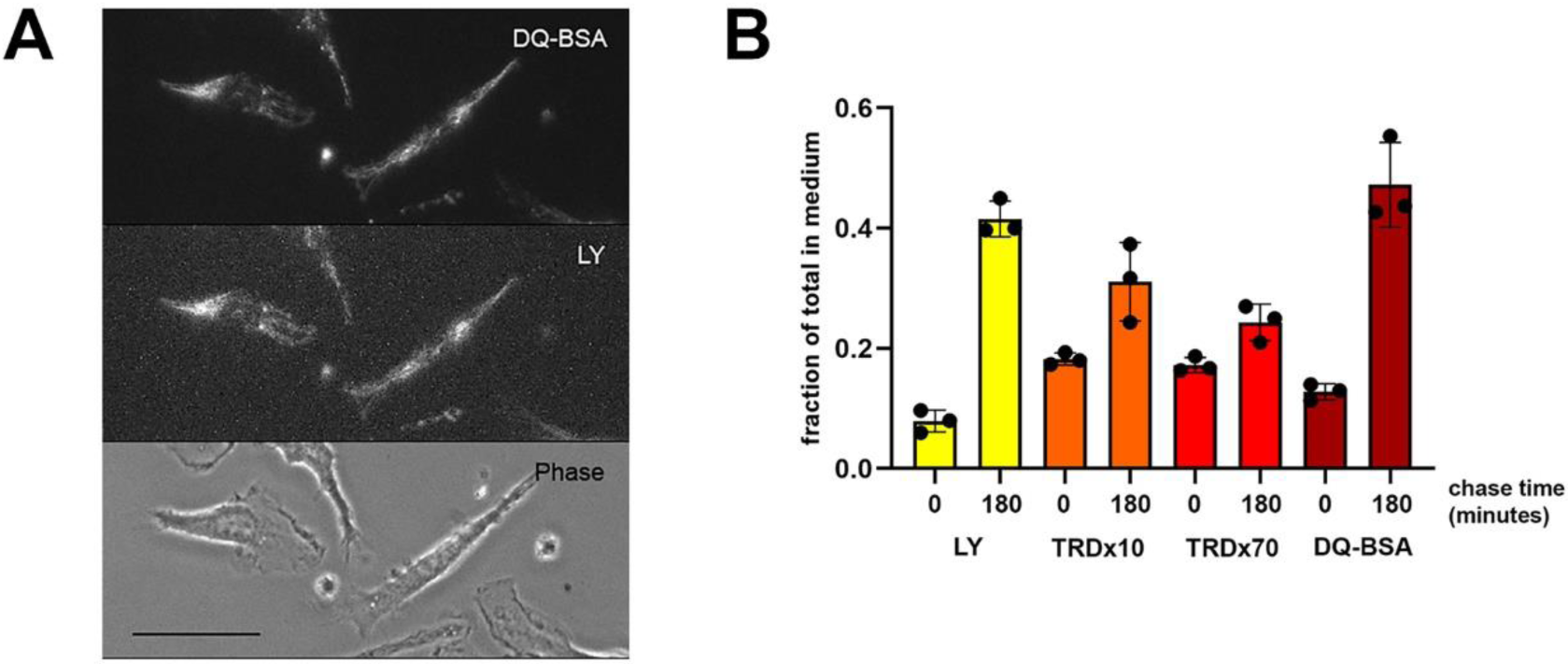
(A) Microscopy of macrophages labeled for 2 hrs with 10 mg/ml DQ-BSA and 500 mg/ml LY in BMM, washed and imaged in RB. Scale bar: 20 mm. (B) Related to the experiments of Fig. 6A, B, showing the fraction of the total fluorescence in each well that was in the medium (vs. cell-associated). The 0-minute chase data were subtracted from the 180-minute chase data to obtain the values in Fig. 6B.

## Notes

### Competing Interest Statement

The authors have declared no competing interest.

